# Phase discontinuities underlie increased drowsiness and diminished sleep quality in older humans

**DOI:** 10.1101/696658

**Authors:** Teresa Hinkle Sanders

## Abstract

Healthy humans switch seamlessly between activity states, wake up and fall asleep with regularity, and cycle through sleep stages necessary for restored homeostasis and memory consolidation each night. This study tested the hypothesis that such smooth behavioral transitions are accompanied by smooth transitions between stable neural states within the brain. A method for detecting phase discontinuities across a broad range of frequencies was created to quantify phase disruptions in the Fp-Cz EEG channel from 20 annotated sleep files. Phase discontinuities decreased with increasingly deep sleep, and increased phase discontinuity was associated with increased drowsiness, reduced deep sleep, and shorter REM sleep. A 10s phase discontinuity summary measure (the *phase jump indicator*) closely tracked the annotated sleep stages and enabled discrimination between short (< 10 min) and longer REM periods. Overall phase discontinuity correlated inversely with broadband EEG power, suggesting that reduced spurious signaling may facilitate increased synchronization. However, the correlation between phase discontinuity and power varied with sleep stage and age. Older individuals spent significantly more time in the Awake and Drowsy stages and less time in the deepest sleep stage and REM sleep. Interestingly, although EEG power was reduced in older individuals across all sleep stages, increased phase discontinuity only occurred in stages that showed impairment. In older patients the power vs. phase discontinuity correlation shifted to positive during drowsiness, suggesting potential deficits in cortical inhibition. These results provide evidence that phase discontinuity measures extend current capabilities for assessing sleep and may yield new insights into pathological brain states.

**Significance statement:** Evidence continues to accumulate regarding the positive relationship between healthy sleep and brain function. Recent studies also show that more healthful sleep can be induced with timely application of non-invasive therapies. Accordingly, the ability to accurately assess sleep quality in real-time has become increasingly important. Here, a newly defined measure, referred to as phase discontinuity, enabled rapid identification of unhealthful neural patterns associated with increased drowsiness, reduced deep sleep, and early termination of REM sleep. Moreover, the measure was linked to underlying neuronal and circuit properties known to impact sleep quality. Thus, the phase discontinuity measure defined in this study provides new insight into sleep pathology and has potential implications for closed-loop therapeutic intervention.

## Introduction

Are there objective neural signatures of healthy brain activity during sleep? Increasing evidence suggests that measures such as power, frequency, and phase-amplitude coupling are useful for assessing sleep. Indeed, EEG recordings can now be used to identify sleep stages with high accuracy (Azim et al., 2010, Lajnef et al., 2015, Dimitriadis et al., 2018), but is there a unifying measure underlying the variations in power and pattern?

### Characteristics of normal healthy sleep stages

During awake states, EEG signals are typically approximately 30 microvolts in amplitude and consist of mixed frequencies with prominent beta and gamma activity (> 15 Hz). Drowsiness, or non-rapid eye movement (NREM) sleep stage 1, is marked by a more variable signal with reduced beta and gamma power. Entry into NREM stage 2 can be identified by increased delta activity (< 4 Hz) and intermittent higher frequency spindles (12-16 Hz). K complexes, sharp negative waves followed by a high voltage slow wave, also appear in Stage 2. In Stages 3 and 4, neuronal activity becomes increasingly dominated by the delta (or slow) wave band. Slow wave activity (SWA) reflects highly synchronized alternating phases of joint hyperpolarization (down states) and depolarization (up states) in large neuron populations. The activity typically reflects traveling waves throughout the cortex, hippocampus, and thalamus. SWA has been reported to enable temporal grouping and synchronization of spindles (and, in turn, bundling of hippocampal ripples), and thus is believed to facilitate communication between these regions (Mölle et al., 2006, Staresina et al., 2015, Varela et al., 2019). This large scale synchronization results in EEG amplitudes that often exceed 75 microvolts. Studies have shown that increased SWA amplitude occurs after sleep deprivation, and that SWA amplitude is restored to nominal levels after sleep (Dijk, 2009, Achermann et al., 1993). Strong SWA during sleep has been linked to enhanced memory performance and boosting SWA during sleep has been shown to improve memory (Bellesi et al., 2014, Ngo et al., 2013). Increased hippocampal sharp wave ripple activity has also been associated with enhanced memory (Fernández-Ruiz et al., 2019).

In normal sleep, rapid eye movement (REM) sleep follows an approximately 90 min period of NREM sleep (Brown et al., 2010). The first REM stage usually exceeds 10 min in length. Four additional REM periods will often occur within a normal night’s sleep (Purves et al., 2001). The REM EEG is a mixed frequency signal characterized by more nominal power levels with a spectral profile that is often similar to that of the Awake and Drowsy stages. Oxygen consumption is decreased during NREM, but increased during REM to levels nearer those typical of wakefulness.

### Age effects

Synapses are pruned during adolescence resulting in reduced delta and theta EEG power (Frank et al., 2001). Delta power (SWA) continues to decrease throughout adulthood (Carrier et al., 2001). Possibly due to their dependence on SWA, sleep spindle intensities and densities have also been shown to decrease with age (Martin et al., 2013).

Additionally, the amount of REM sleep decreases with age. Infants often experience 8 hours of REM sleep per 24 hours. By age 20 the typical amount of REM sleep is approximately 2 hours per night, while at age 70, the average is reduced to 45 min (Purves et al., 2001). Conversely, sleep fragmentation and time spent in awake and drowsy stages each night increase with age (Bonnet, 1989). Although many older individuals maintain healthful sleep and high cognitive function (Anderson and Horne, 2003, Huber et al., 2004, Carrier et al., 2001), sleep pathology typically increases with age and has been shown to correlate with decreased performance on memory tasks (Van Someren et al., 2015, Mander et al., 2017). Given the important relationships between sleep and brain clearance, memory, cognition, and psychological health (Xie et al., 2013, Sara, 2017, Brayet et al., 2016, Anderson and Horne, 2003), new measures are needed that provide early identification of pathological neural signaling related to disrupted sleep.

This study examines whether measurements of phase discontinuity are useful for understanding sleep progression in younger and older individuals, and how these measures relate to current spectral power benchmarks used to evaluate sleep. Further, the study explores whether phase discontinuity may enable detection of pathologies related to cortical inhibition and neuronal synchronization. The results show that EEG measures of phase discontinuity as defined in this study extend current capabilities for assessing sleep quality and may provide important early insight into an individual’s capacity for healthy brain function.

## Results

To test whether phase discontinuities play an important role in wake/sleep stages, 20 sleep data files recorded from 10 subjects were analyzed (Table 1). Six wake/sleep stages were examined: Awake, Drowsy, Non-Rapid-Eye Movement sleep stages 2 through 4 (NREM2-4), and Rapid-Eye-Movement (REM) sleep. Time-frequency phase difference plots and a summary measure of phase discontinuity (*phase jump indicator*, see Methods), were used to quantify the phase change activity over time.

**Table 1.** Number of 10 s epochs in sleep stages for each sleep file, ordered by age.

| Age | Sex | Awake | Drowsy | NREM2 | NREM3<br>+<br>NREM4 | Total<br>REM | 1st<br>REM<br>period | avg.<br>REM<br>period | num.<br>REM | num.<br>REM<br>< 10<br>min |
| --- | --- | --- | --- | --- | --- | --- | --- | --- | --- | --- |
| 20 | F | 375 | 216 | 909 | 915 | 693 | 53 | 86.625 | 8 | 3 |
| 20 | F | 159 | 90 | 774 | 882 | 1029 | 143 | 205.8 | 5 | 0 |
| 21 | F | 96 | 111 | 1164 | 633 | 879 | 152 | 175.8 | 5 | 0 |
| 21 | F | 90 | 75 | 1224 | 672 | 945 | 140 | 157.5 | 6 | 1 |
| 21 | M | 177 | 102 | 1356 | 609 | 801 | 110 | 160.2 | 5 | 0 |
| 21 | M | 264 | 219 | 1440 | 474 | 432 | 50 | 86.4 | 5 | 1 |
| 24 | M | 42 | 195 | 1830 | 276 | 402 | 11 | 44.7 | 9 | 8 |
| 24 | M | 45 | 102 | 1875 | 132 | 522 | 14 | 47.5 | 11 | 9 |
| 28 | F | 42 | 63 | 1635 | 336 | 696 | 56 | 99.4 | 7 | 3 |
| 28 | F | 204 | 204 | 1269 | 579 | 600 | 89 | 100 | 6 | 1 |
| 35 | F | 111 | 267 | 1767 | 30 | 813 | 53 | 101.6 | 8 | 2 |
| 35 | F | 111 | 141 | 1743 | 78 | 738 | 77 | 52.7 | 14 | 10 |
| 48 | F | 54 | 114 | 2010 | 135 | 711 | 68 | 71.1 | 10 | 4 |
| 48 | F | 42 | 66 | 1773 | 417 | 510 | 71 | 102 | 5 | 1 |
| 56 | M | 624 | 633 | 1314 | 6 | 693 | 140 | 46.2 | 15 | 10 |
| 56 | M | 234 | 573 | 1425 | 0 | 600 | 86 | 30 | 20 | 16 |
| 66 | F | 423 | 513 | 1350 | 135 | 333 | 98 | 111 | 3 | 0 |
| 66 | F | 462 | 237 | 1404 | 504 | 264 | 98 | 66 | 4 | 0 |
| 79 | F | 456 | 306 | 1290 | 654 | 240 | 20 | 26.7 | 9 | 5 |
| 79 | F | 171 | 276 | 1335 | 612 | 304 | 104 | 152 | 2 | 0 |

### Phase discontinuities decrease with transitions to deeper sleep (NREM2-NREM4)

During Awake and Drowsy stages the EEG phase profile was dynamic with wide swaths of phase discontinuities throughout time-frequency difference space (Fig. 1a). However, as NREM sleep progressed, the phase profile and *phase jump indicator* reflected increasingly reduced phase change activity (Fig. 1b and 1f). Quantification of the distribution of phase changes > |π/5| confirmed the largest amount of phase discontinuity occurred in Awake (Fig. 1c) and Drowsy states (mean(*phase jump indicator)* = 0.28, 0.24 respectively, p = 1.6e-43) with significantly reduced phase discontinuity in successive NREM stages (mean(*phase jump indicator)* = 0.11, 0.05, 0.03 for NREM2-4, p < 6.5e-159, Fig. 1d and 1e). The phase discontinuity distribution for each sleep stage was significantly different from all other stages (p < 2e-43, Fig. 1d).

**Figure 1.**
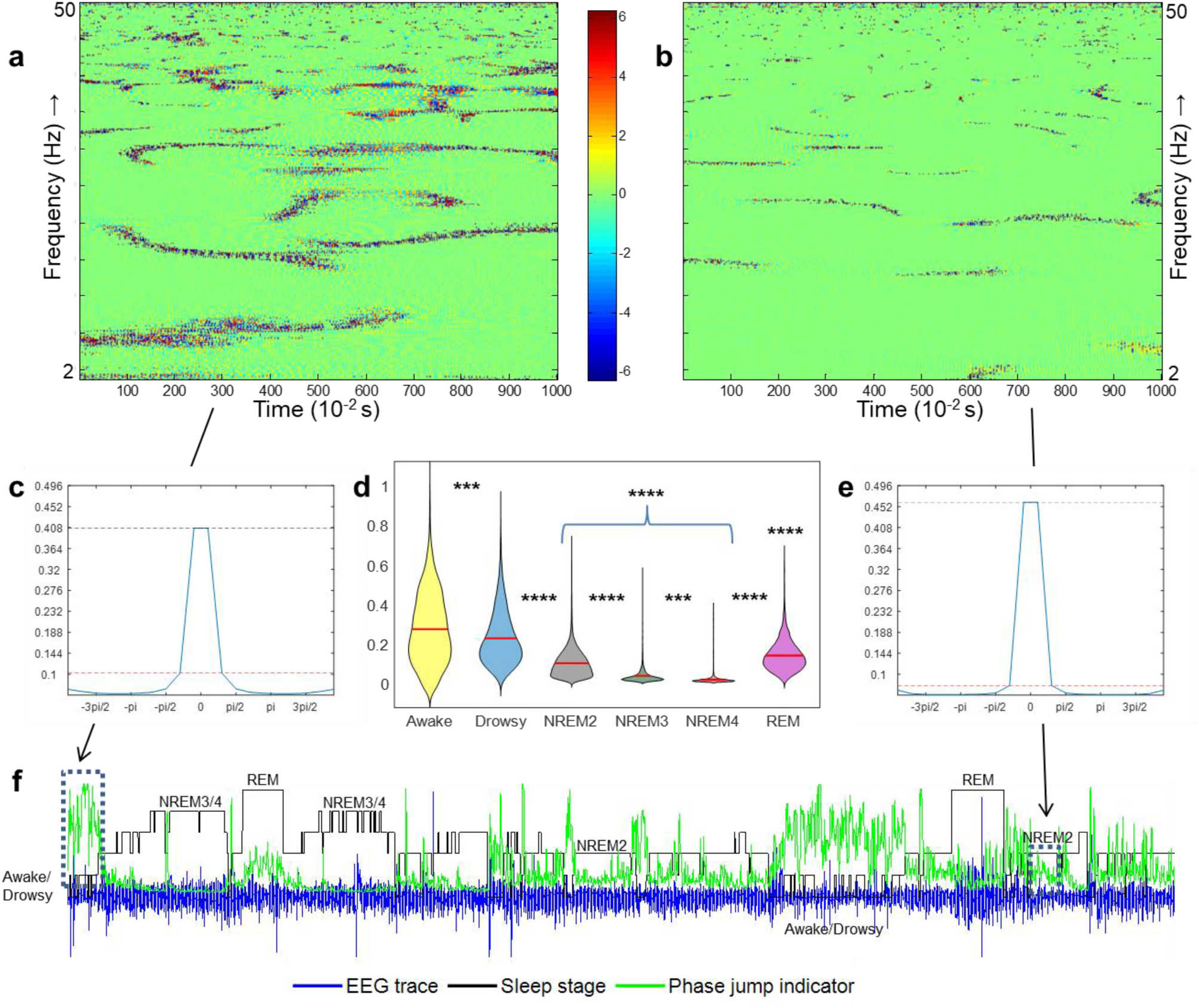
The degree of phase discontinuity varies by sleep stage. Continuous wavelet transform time-frequency difference plots of representative 10 s EEG traces during **(a)** Awake and **(b)** NREM2 sleep stages. **(c)** Normalized histogram of all the Awake time-frequency difference plots. **(d)** *Phase jump indicator* by sleep stage for all study subjects combined. **(e)** Normalized histogram of all the NREM2 time-frequency difference plots. **(f)** EEG trace (blue), expertly scored sleep stages (black), and *phase jump indicator* (green) for subject 16 (female aged 79). **** p < 1e-158, *** p < 1e-42 upper row of asterisks reflects significance between Awake and other stages, lower row of asterisks reflects significance between adjacent sleep stages.

### REM sleep stage length can be classified by examining phase discontinuities

Previous studies have found that healthy initial REM stages are typically at least 10 min in length (Purves et al, 2001). In the current study, examination of REM sleep stages by length revealed that, although REM sleep showed a moderate average level of phase discontinuity (0.13) (Fig 1d), shorter REM stages had significantly higher phase discontinuity and more variability (Fig. 2a-c). *Phase jump indicators* over five consecutive 10 s epochs enabled ∼ 80% accuracy in distinguishing between shorter (< 10 min) and longer (≥ 10 min) REM stages (performance with *phase jump indicator* alone = 78% (vector length 6), with power band measures = 82% (vector length 36), boot strap aggregation results, number of trees = 100, 5-fold cross validation, Fig. 2d).

**Figure 2.**
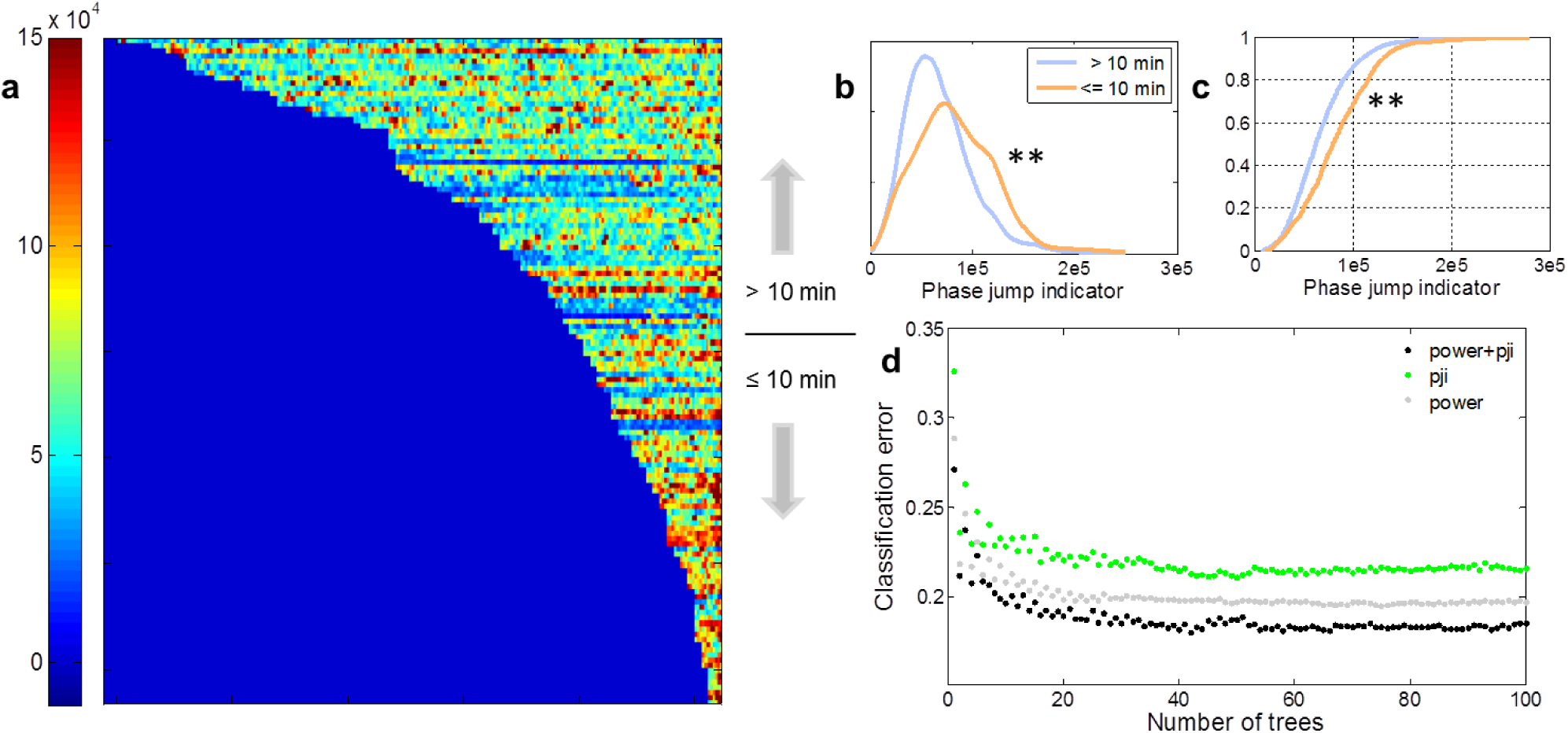
Phase discontinuity measures enable discrimination between short and long REM sleep periods. **(a)** Rows show the 133 REM stages from all subjects, sorted by length. Pixels within each row show the intensity of the *phase jump indicator* for each 10 s epoch in each REM period. **(b)** The *phase jump indicator* density for REM stages ≤10 min was broader and shifted upward (orange) compared to the density for REM stages > 10 min (light blue), **(c)** Cumulative distribution for the data shown in (b). **(d)** Vectors of 5 consecutive *phase jump indicators* enabled discrimination between short (< 10 min) and long (> 10 min) REM stages. Combining *phase jump indicators* and spectral power measures enabled increased discrimination but required a much longer feature vector (feature vector length = 36). ** p < 1e-4. pji = phase jump indicator

### Phase discontinuity correlation with power varies by sleep stage

The band power profiles associated with different sleep stages and their relationship to neural synchronization are well documented (Brown et al., 2010, Brayet et al., 2016, Carrier et al., 2001). In this study, to test a secondary hypothesis that phase discontinuity might disrupt synchrony and thus be inversely related to EEG power, the relationship between phase discontinuity and power was analyzed (Fig. 3). Pearson correlation results revealed that, over all wake/sleep stages combined, the *phase jump indicator (pji)* indeed correlated negatively with power (r_pji,p_ = **−** 0.22, p < 1.0e-274). Notably, however, the correlation between *phase jump indicator* and power, r_pji,p_, changed from negative during wakefulness (r_pji,p_ = **−** 0.20, p = 2.8e-36), to slightly positive during drowsiness (r_pji,p_ = +0.05, p = 9.6e-4). As expected due to the high degree of neural synchrony during NREM2-4, phase discontinuity again correlated negatively with power during these stages (r_pji,p_ = −0.15, −0.12, −0.21, respectively, p < 1.3e-13, Fig. 3b). Interestingly, the strongest negative correlation occurred during REM sleep (r_pji,p_ = **−** 0.31, p = 1.8e-274). This overall negative correlation was supported by negative correlations with power in the delta and theta bands (r_pji,p_ = −0.47, −0.45 respectively, p < 1.0e-42) during REM sleep.

**Figure 3.**
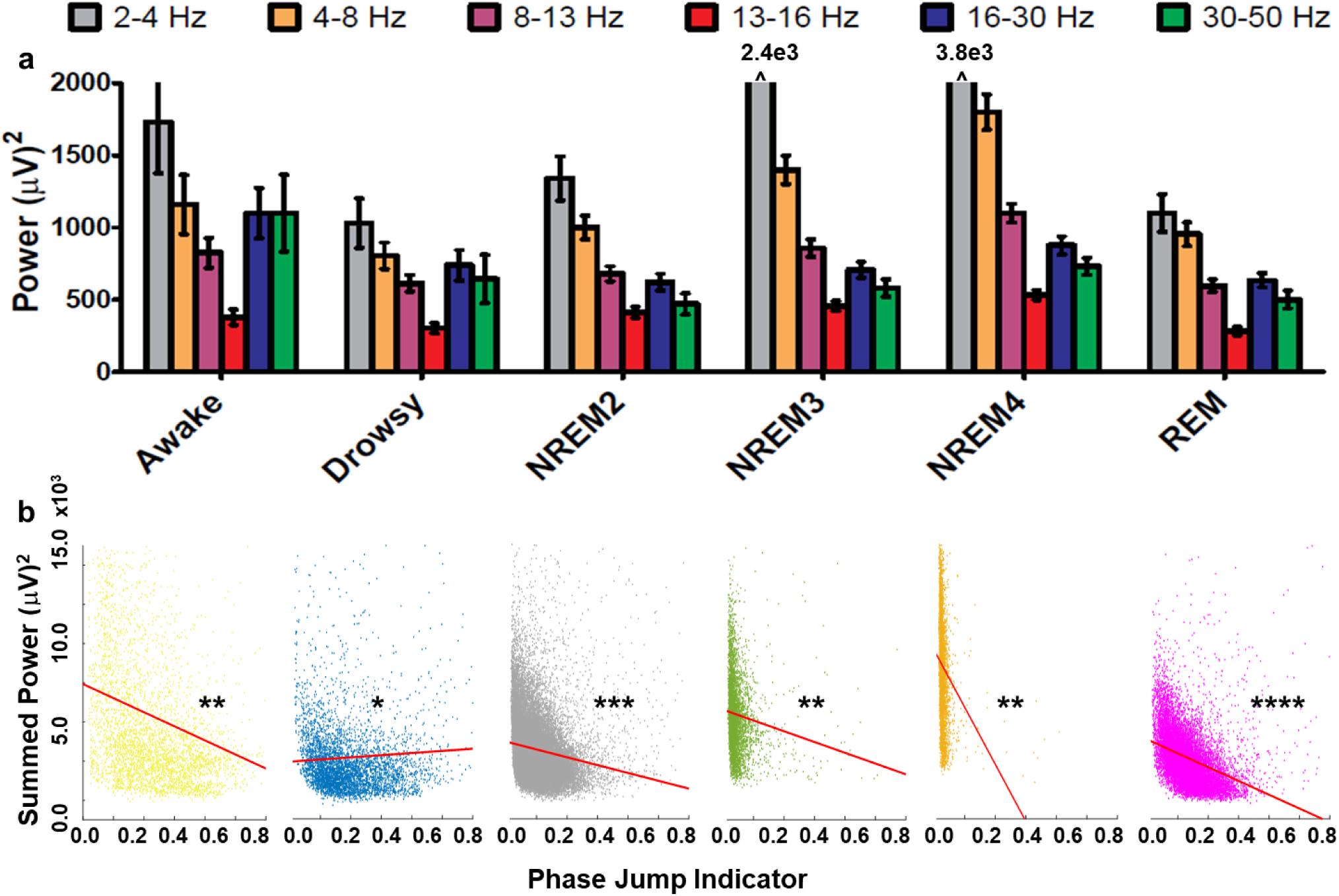
Power and phase discontinuity vs. power correlation varies by sleep stage. **(a)** Spectral band power for all subjects combined, and **(b)** Scatterplots of the *phase jump indicator* vs. total power for all subjects. Red lines indicate least squares fit for each scatterplot. Note that although the spectral power profiles for Drowsy and REM stages are similar, the correlation between power and *phase jump indicator* for the Drowsy stage was positive (r = +0.05), while the REM stage correlation was negative (r = −0.31). **** p < 1e-158, *** p < 1e-42, **p < 1e-4, * p < 0.05. Error bars reflect SEM.

### Older subjects displayed more awake/drowsy time, and shorter periods of deep (NREM4) and REM sleep

Consistent with previous sleep studies (Carrier et al., 2001, Mander et al., 2017), age and time in Awake and Drowsy stages were positively correlated (Figs. 4a, Awake: r_A,a_ = 0.52, p = 0.02, Drowsy: r_A,d_ = 0.58, p = 0.01). Age was negatively correlated with time in the deepest sleep stage and REM sleep (NREM4: r_A,N4_ = - 0.43, p = 0.05, REM: r_A,R_ = - 0.59, p = 0.01). The length of the first REM stage and the total time in REM sleep were also correlated (r_REM,REM1_ = 0.57, p = 0.008), highlighting the importance of the first REM stage to overall sleep quality (Table 1, average number of REM stages = 7.85, no other REM stage correlated with total REM stage time). In the 20 sleep files examined, 1/3 of the first REM periods were < 10 min (below the healthy standard).

**Figure 4.**
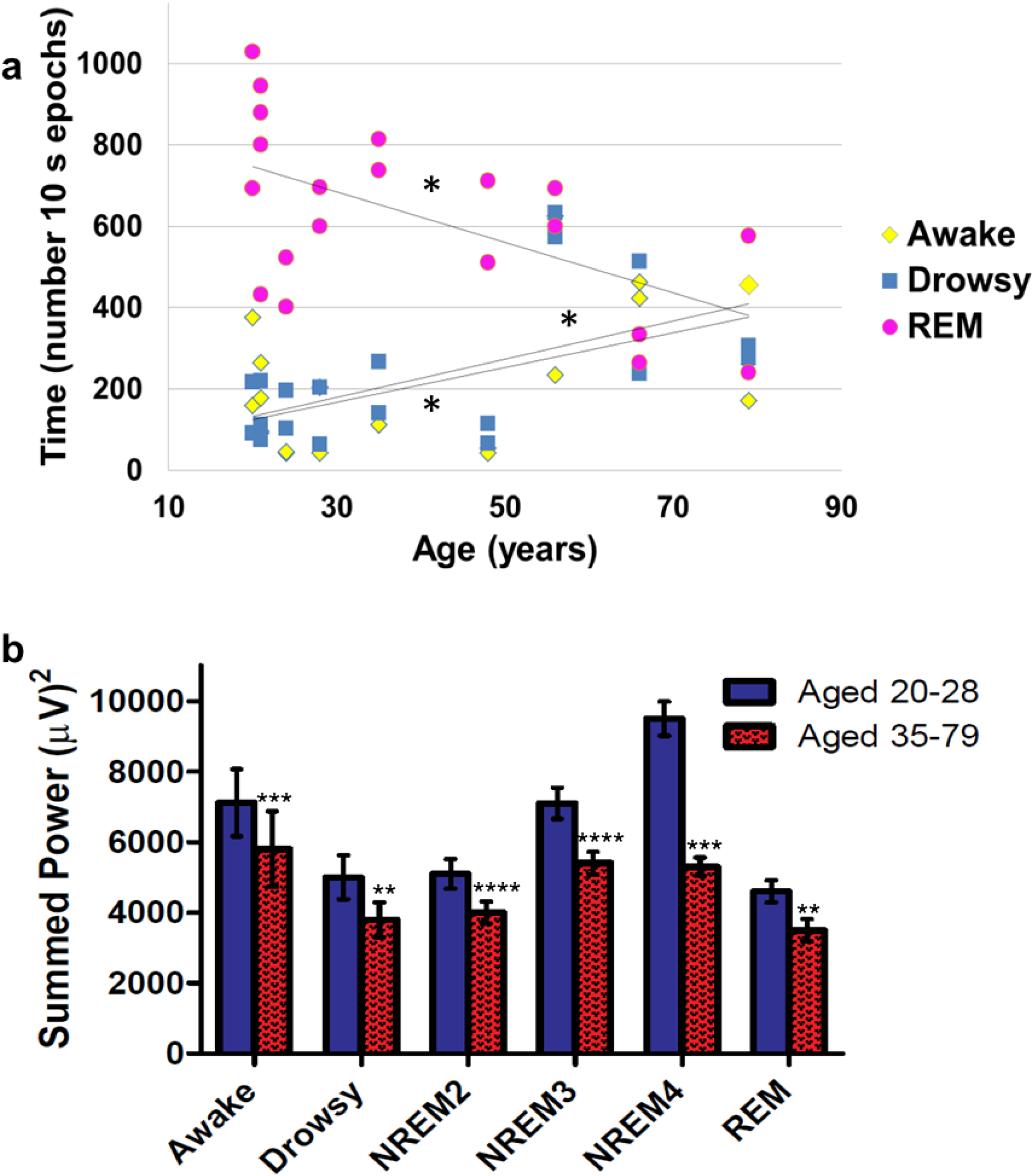
Age-related changes in waking and sleep. **(a)** Time spent in Awake and Drowsy stages generally increased with age, while time spent in REM stage decreased with age. **(b)** Average broadband power (2-50 Hz) decreased in older subjects. **** p < 1e-158, *** p < 1e-42, ** p < 1e-4, * p < 0.05

Grouping the subjects into two equal sub-populations by age (population 1: 20-28, population 2: 35-79) revealed that older subjects spent over twice as many epochs in the Drowsy stage compared to younger subjects (3126 / 1493 10 s epochs, older / younger). Younger subjects spent over 4.8 times as many epochs in NREM4 compared to the older subjects (3288 / 681 10 s epochs, younger / older).

### Older subjects displayed reduced total power, overall increased phase discontinuity, and significantly changed correlation between power and phase discontinuity

Comparison of the EEG data grouped into older and younger sub-populations revealed reduced power (overall reduction ratio: 0.7, p < 1.0e-274, Fig. 4b) and overall increased phase discontinuity (ratio: 1.1, p = 3.1e-88, Fig. 5a) in the older vs. younger groups. Examination by sleep stage revealed that the mean *phase jump indicator* was increased in the older vs. younger groups in Awake (1.3, p = 8.4e-47), Drowsy (1.1, p = 3.3e-11), NREM4 (1.1, p = 6.7e-6), and REM (1.2, p = 5.8e-118) sleep, but was reduced in NREM2 (0.8, p = 7.9e-220) and NREM3 (0.7, p = 7.1e-25). Interestingly, the stages indicating increased phase discontinuity were also the stages that showed impairment with age (increased Awake and Drowsy time, reduced NREM4 and REM time). Notably, despite a decreasing trend in phase discontinuity in both older and younger subjects during NREM2 and NREM3, the rate of change between NREM stages was smaller in older subjects (−2.6e4 vs. −3.1e4), such that by the time NREM4 was reached, the older group again had increased phase discontinuity relative to the younger group.

**Figure 5.**
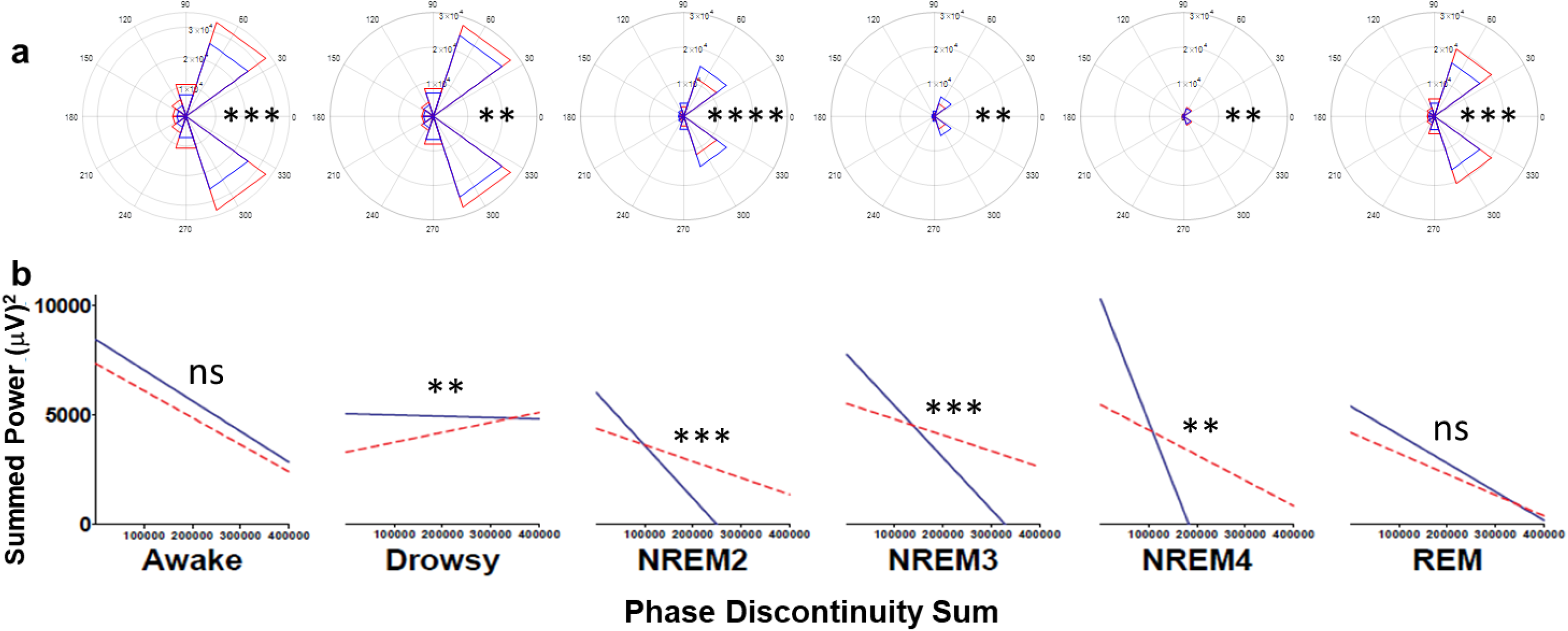
Age-related changes in phase discontinuities and correlation with power. **(a)** Phase discontinuities were increased in the older group (red) in Awake, Drowsy, NREM4, and REM sleep, but were reduced in NREM2 and NREM3 compared to the younger group (blue). **(b)** Correlation between total power and phase discontinuity was not significantly changed in the older (red dashed line) vs. younger (blue solid line) subjects during Awake and REM stages, but differed significantly in Drowsy and NREM stages. Note that the range of variation in slope between NREM sleep stages was reduced in older vs. younger subjects. **** p < 1e-158, *** p < 1e-42, **p < 1e-4

The overall correlation between the *phase jump indicator* and power also varied significantly between the two age groups (Pearson correlation, r_pji,p_ = −0.37 vs. −0.07 in the younger vs. older populations, p < 1e-5). Examination of these correlations by sleep stage revealed that, while the r_pji,p_ correlations were not significantly changed during awake periods (Fig. 5b, −0.17 for both younger and older) and REM sleep (−0.27 vs. −0.25 for younger and older, respectively), the correlations during Drowsy and NREM sleep stages were significantly different (−0.01 vs. +0.12 during drowsiness, p = 3.4e-5, and {−0.27, −0.27, −0.24} vs. {−0.17, −0.09, −0.11} for NREM2 through NREM4 in younger vs. older, p < 1e-3).

## Discussion

Brains attend to change and brain circuitry is adapted to detect alterations in the electrophysiological waves generated by populations of neurons. This biological framework and the mathematical relationship between phase discontinuities and altered waveforms led to the hypothesis that excessive fluctuations in phase may reduce brain efficiency and disrupt brain state transitions. Clearly, some discontinuities are necessary for switching activity patterns and states in dynamic environments, but in the healthy brain these discontinuities are typically brief and transient, e.g., beta bursts in movement and shifting spectral profiles for sleep stage transitions. In pathological conditions, where waveform amplitudes and shapes may be driven by random spiking due to either increased biological noise or reduced signal and disrupted circuitry due to cell death, discontinuities may be recurring and waveform irregularities may be prolonged.

### Physiological basis of reduced phase discontinuity in sleep

The *phase jump indicator (pji)* summary measure created in this study to examine phase discontinuities captures the progression from waking to deeper sleep stages remarkably well, in agreement with the intuitive reasoning that disruptive phase discontinuities would be reduced during the transition to deeper sleep. These results also support the hypothesis that healthy behavioral transitions accompany smooth neural transitions, since healthful sleep transitions to deeper sleep states occurred in tandem with reduced phase discontinuity.

However, while reduced phase discontinuity was clearly apparent in the transitions to deeper sleep, single *pji* measures within REM sleep were easily confused with measures from other stages. Further analysis of the measures in the context of REM sleep stage length revealed that the *pji* mass function was right-shifted and broader in short REM stages (≤ 10 min), suggesting that reducing both phase discontinuity and variability is important for maintaining longer REM sleep stages.

During sleep, when the cortex is largely disengaged from external inputs, phase discontinuities may represent noise or internal pathologies in the brain. The inability to maintain longer duration REM stages and the deepest sleep stage (NREM4) observed in some patients in this study may be due to an internal ‘noise floor’ in the brain. The observed increased phase discontinuity and variability during short REM stages suggest these measures may be useful for estimating the noise floor.

### The relationship between phase discontinuity and power

The finding of overall negative correlation between phase discontinuity and power supported this study’s secondary hypothesis that reduced phase discontinuity is important for increased signal power. This suggests that, in order to promote healthful sleep, it is important for the brain to reach a quiescent state where sensory stimuli are attenuated and phase discontinuities limited to those essential to propagate the population dynamics required for the nascent neural processing. Sensory stimuli that are not adequately suppressed by cortical interneurons, and internal sources of noise, such as inflammation, may induce random phase discontinuities that reduce the ability of the brain to maximize the amplitude of the oscillations necessary for memory consolidation and other important processes that occur during sleep.

Surprisingly, the detailed Pearson correlation analysis revealed that the correlation between phase discontinuity and power was highly stage specific. In particular, the Drowsy stage correlation was slightly positive while all other *pji*-power correlation measures were negative. This unique relationship between phase and power during drowsiness may be related to the disengagement of external sensory inputs during the transition from waking to sleep. In other words, as measured in this study, EEG phase discontinuities during the Awake stage are likely driven by large shifts in activity, sensory input, or other global state changes. Interneurons and cortical networks both respond to and adapt to fold these inputs into ongoing neural activity so that signal power and structure is preserved and optimized. In order to transition to sleep, these sensory inputs are believed to be disengaged through inhibition of thalamacortical neurons via GABAergic activity in the reticular thalamus (Steriade, 2005). At the same time, cortical neurons begin to synchronize for SWA. When this transition occurs, it is likely that the sensory-driven phase discontinuities may persist until the disengagement is stable and the new internally-driven cortical oscillatory pattern emerges. At this point, the externally-driven discontinuities may no longer be involved in meaningful cortical signaling, thus the relationship between these inputs and signal power becomes close to random.

Disengagement between phase discontinuity and EEG broadband power in healthy Drowsy stages is supported by the almost entirely random relationship between the two in the younger subjects in this study. In contrast, the marked increase in the Drowsy *pji*-power correlation slope in older patients suggest that, rather than being suppressed, the discontinuities may actually be driving the population signal. The pathological nature of this signaling activity is further suggested by the significantly increased time that older patients spent in the Drowsy stage.

### Phase discontinuity as a biomarker of pathophysiology

The relationship between EEG power and phase discontinuity discovered in this study indicates that, although the younger and older subjects maintain a similar degree of correlation between power and phase discontinuity during the Awake stage, the correlation is significantly altered during abnormal NREM stages observed in older patients. The reduced magnitude of the *pji*-power negative correlation during deeper sleep stages may reflect an inability to synchronize neurons due to internal noise, insufficient numbers of healthy neurons, or circuit maladaptation. It is also possible that the greater power displayed by the younger subjects is related to a larger inherent dynamic range that enables more adaptability to input discontinuities (steeper slopes in the *pji*-power graphs). Alternatively, the higher power signals in younger subjects may increase the signal-to-noise ratio (SNR), thus reducing the effect of noise-driven phase discontinuities. Regardless of the underlying mechanism, the shallower slopes and similar appearance of the NREM *pji*-power graphs in older subjects suggest reduced flexibility and dynamic range. Thus, it may be useful to view the *pji*-power negative correlation as a measure of brain efficiency with regard to translating individual neural signals within a population into the high power, smoothly varying oscillations required for effective sleep. Note that the significant phase discontinuity vs. power correlation changes that occur in older subjects during sleep stages but not during wakefulness, support findings from other studies that sleep analysis is useful for detecting early brain pathophysiology (Lucey et al., 2019, Stiasny-Kolster et al., 2005).

To summarize, in simplest terms this study is an analysis of signal vs. noise in biological channels. The results point to the close intertwining of signal, noise, and channel in the brain, and enable quantification of the degree to which the population signal and diverging inputs from subgroups of cells are altered and (inversely) correlated during different brain states, thus providing indirect measures of the health and efficiency of the underlying cells and their connectivity. Specifically, the new CWT time-frequency phase discontinuity plots, *phase jump indicator*, and correlation between phase discontinuity and power presented in this study identify top-level EEG signal characteristics important for deeper sleep stages, longer REM sleep stages, shorter periods of drowsiness, and overall healthful sleep patterns. Thus, these measures are useful biomarkers for identification of sleep pathologies. Additionally, the high update rate, easy sleep state readout, and minimal computation associated with the *phase jump indicator* makes this summary measure of interest for real-time EEG analysis and potential feedback in adaptive neural modulation strategies. Future Deep Learning analyses of the time-frequency phase discontinuity plots in multi-recording site data with a higher sampling rate (> 100 Hz) from a larger group of subjects will be of interest. Such analyses may reveal the impact of important signaling constructs such as sharp wave ripples on the top-level population signal structure examined in this study.

## Methods

Analysis was performed on 20 sleep recordings from 10 individuals (males and females) in the set of Physionet Sleep-EDF Database “ST” files (Kemp et al., 1993, Kemp et al., 2000, Goldberger et al., 2000) previously used by the author to evaluate automatic Sleep Stage Classification (Sanders et al., 2014). Study subjects had some difficulty falling asleep but were otherwise healthy. Recordings from the Fpz-Cz EEG electrodes were examined in both baseline and medicated (Temazepam) conditions. Data from the two nights were pooled since no significant differences in sleep quality were found between the medicated and unmedicated conditions. Each of the 20 files analyzed in this study contains approximately 8 hours of sleep. Sleep scoring annotations (hypnograms) were available for each file: Movement, Waking, Rapid Eye Movement (REM) sleep, and non-REM sleep stages 1-4 (Drowsy = NREM1).

Phase discontinuities were calculated from the 100 Hz EEG as follows. The complex continuous wavelet transform (CWT) was applied using the following formulation to obtain changes in the phase of the coherence between consecutive 10 s epochs across 512 wavelet scales corresponding to frequencies from 2 Hz to 50 Hz at each time sample (time-frequency phase plots).

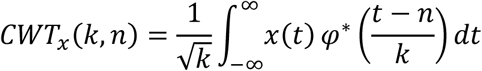

Where “k” represents the scale factors, “n” represents the time points over which the CWTs are evaluated, and *φ*(t) is the complex Morlet wavelet:

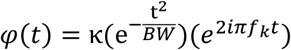

The coherence between each adjacent 10 s epoch was calculated, and the time-frequency phase plot was obtained.

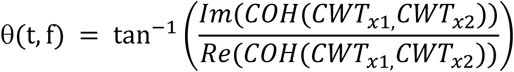

Consecutive time points in the time-frequency phase plots were subtracted to identify phase changes occurring over short time intervals (difference plots, Figs. 1a and 1b). The phase changes from the difference plots were histogrammed into 20 bins ranging from −2π to +2π (Figs. 1c and 1e), denoted pdf(Δθ_i_). The summary *phase jump indicator, pji*, measure was then defined as the ratio between the total number of phase “jumps”, Δθ > |π /5|, and smaller phase changes, Δθ ∈ [-π /5, + π /5] (Figs. 1d and 1f).

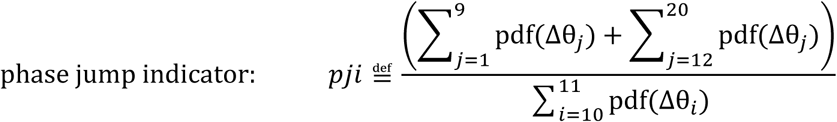

The frequency band power was calculated using the method described in the author’s previous sleep classification study (Sanders et al., 2014). Briefly, data was segmented into 1 second frames and the power within each of the Delta (3-4 Hz), Theta (4-8 Hz), Mu/Alpha1 (8-13 Hz), Alpha2/Beta1 (13-16 Hz), Beta2 (16-30 Hz), and Gamma (30-50 Hz) frequency bands was calculated using short-time Fourier transforms (STFTs). For comparison, the power within each band for each 10-sec epoch was averaged. Note that, although the bands are the same as those used in the previous study, they are slightly re-named here to reflect frequencies more commonly used for sleep study.

For REM length classification, the performance of feature vectors containing *phase jump indicators* for each 5 consecutive 10 s epochs along with the *phase jump indicator* variance across the 5 epochs (feature vector length = 6) was compared to the performance of these 6 features combined with spectral power features from the 6 above frequency bands over the 5 epochs (feature vector length = 6 + 6×5 = 36). The corresponding labels from the annotation files were used to enable classification with 5-fold cross-validation using the bootstrap aggregating (bagging) algorithm method described in the previous study (Sanders et al., 2014).

Since differences in power between male and female EEGs have previously been reported (Carrier et al., 2001), the data were examined for sex effects. Male subjects (2 in the younger group and 1 in the older group) were removed and the top-level measures between the female-only and pooled sex cohorts were compared to ensure the results were not significantly changed by including both sexes.

General significance was assessed with 2-sided, 2 sample t-tests. Correlation and significance between age and sleep stage statistics, and between phase discontinuity and power measures were evaluated using Pearson correlation (MATLAB *corrcoef*). Comparisons between correlations (r) were evaluated by performing the Fisher’s r to z transformation, then analyzing the significance of the observed z test statistic (α = 0.05).

## Notes

#### Summary of Updates

The abstract has been updated to clarify the important findings.

https://physionet.org/physiobank/database/sleep-edfx/

